# *sigQC*: A procedural approach for standardising the evaluation of gene signatures

**DOI:** 10.1101/203729

**Authors:** Andrew Dhawan, Alessandro Barberis, Wei-Chen Cheng, Enric Domingo, Catharine West, Tim Maughan, Jacob G. Scott, Adrian L. Harris, Francesca M. Buffa

## Abstract

With the increase in next generation sequencing generating large amounts of genomic data, gene expression signatures are becoming critically important tools, poised to make a large impact on the diagnosis, management and prognosis for a number of diseases. Increasingly, it is becoming necessary to determine whether a gene expression signature may apply to a dataset, but no standard quality control methodology exists. In this work, we introduce the first protocol, implemented in an R package sigQC, enabling a streamlined methodological and standardised approach for the quality control validation of gene signatures on independent data sets. The emphasis in this work is in showing the critical quality control steps involved in the generation of a clinically and biologically useful, transportable gene signature, including ensuring sufficient expression, variability, and autocorrelation of a signature. We demonstrate the application of the protocol in this work, showing how the outputs created from sigQC may be used for the evaluation of gene signatures on large-scale gene expression data in cancer.

## Introduction

Gene signatures, over the past decade have revolutionised our understanding of disease, pathogenesis, and clinical response [4, 10, 15]. The application of gene signatures to the clinic has become a massive force driving healthcare forwards towards personalised medicine, and in doing so, has led to a large development effort. These gene signatures are derived by an ever-increasing arsenal of methodologies, spanning approaches such as supervised [14] and unsupervised clustering [8], seed-based approaches [3] and other machine-learning techniques [7]. However, in many cases these signatures remain limited by narrow use cases, or display a general ability in predictive power not specific to any particular state of disease, with even random gene signatures capable of significantly separating groups of breast cancer patients with favourable and unfavourable outcomes [17]. In response to these results, Berglund et al. proposed a framework by which signature coherence, uniqueness, robustness, and transferability are evaluated for PCA-based signatures, and show how these may be applied to such signatures as a check for validity [2].

The pertinent application of gene signatures to a vast array of clinical data depends critically upon the ability of the signature to perform robustly over a wide range of possible confounders, noise, and inter-platform differences for gene expression profiling [9]. In order to ensure that the influence of such factors is reduced, we propose, within this work, a battery of tests and validation criteria that, if passed, would ensure that these competing effects are reduced. Specifically, the R package *sigQC* was developed to standardise and simplify the quality control metrics used to evaluate the applicability of a gene signature to a given dataset, summarising a series of established metrics (see e.g. [3] or [11]) whilst deriving compact metagenes. To illustrate this tool in this work, the use case of *sigQC* in evaluating a published gene signature for breast cancer metastasis on clinical tumour samples with gene expression measured by RNA-seq and microarray is shown, in addition to a signature comprised of a random set of genes to highlight the differences in performance between a well-performing signature and a poor-performer with respect to the metrics produced by the *sigQC* tool. We show how the output of the quality control plots change in the presence of a high-performing signature and a relatively poorly performing signature on a particular dataset, and how a signature may be tested to ensure cross-platform applicability.

## Overview of the protocol

Conceptually, this protocol is designed to ensure that gene signatures are derived with characteristics suitable for clinical utility, and to elicit those properties that pertain to broader application. During the process of derivation of a new gene signature or assessment of an existing one, there are many different aspects that must be accounted for: i) signature technical transportability, ii) signature biological integrity, iii) signature suitability and iv) dataset suitability. To clarify further, signature transportability refers to the use of a gene signature across datasets produced by different technologies, such as RNA-seq vs. microarrays, which quantify genes differently, though they may come from the same sample or origin. The importance of this is underscored by the fact that over the previous decade, most gene signatures have been developed using DNA microarray technology, i.e. a collection of DNA ‘probe’ sequences attached to a solid surface, but most sequencing at present is done by RNA-seq. However, this is further complicated by the fact that microarrays themselves comprise a range of technical methodologies (e.g. spotting, in-situ synthesis) and may have different output characteristics (e.g. one-channel vs. two-channel detection) [13]. Moreover, presently, whole transcriptome sequencing (RNA sequencing) is increasing in popularity and decreasing in cost, providing a wealth of genomic data, quantified in yet a different manner, and represents the current trajectory forward in the technological development of new gene signatures [18]. All of this variability between technologies must be taken into account, and the behaviour of a gene signature should show consistency across datasets generated by these technologies.

Secondly, given datasets generated using the same technology, a signature’s ability to represent a biological phenomenon in a general, reproducible sense in a specific context, should be ensured, before moving on to wider application. For instance, gene signatures derived in a general sense for cancer, should first encapsulate all of the heterogeneity in, for instance, breast cancer, before being tested on colon cancer.

Lastly, in the case of multiple signatures and multiple datasets for the same phenotype or biological process being captured, the signature under primary consideration should be the most suitable for both the dataset and the level of generalizability desired. Further, it should be ensured that each dataset possesses properties suggesting signature compatibility before proceeding with further analysis. Once these have been ensured, quality control via this protocol may proceed.

There are two overarching aspects to this protocol, the first being the tests of the properties of the genes comprising the signature itself, and the second being the properties of the dataset as it pertains to the signature genes. A flowchart of the procedure is depicted in Figure 1.

**Figure 1:**
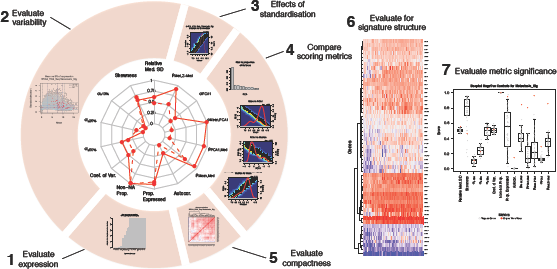
Flowchart of steps involved in the proposed *sigQC* protocol and sample output plots produced by the *sigQC* R package.

The evaluation of the genes composing the signature is primarily to determine whether signature genes are those that co-operate to give a strong, coherent signal across the samples, which are both expressed and varying. Furthermore, the distribution of expression values for signature genes should be coherent enough to be summarised into a robust value for comparison across samples, and these tests are described within this protocol.

As important as the signature itself, are the statistical properties of the dataset to which it is applied. Thus, within this protocol, we describe how a search for structured subcomponents of a gene signature or dataset may be done, to discover whether there are subsets of genes or samples that could benefit from subsetting as a distinct class. Furthermore, the *sigQC* package includes commands for bootstrapping, or the evaluation of a set of negative controls – random gene sets of the same length as the signature itself, to reveal the underlying null distributions of each of the metrics we consider in evaluating signature quality.

## Application of the method

Here, we depict a motivating example of the protocol, as implemented through the R package *sigQC* to evaluate a published gene signature for breast cancer metastasis, and a random gene signature on a RNA-seq dataset from clinical breast tumour samples, downloaded through the Firebrowse portal as part of the Cancer Genome Atlas project (TCGA) [1, 12]. The gene signature used is a set of upregulated genes in breast cancer metastasis, taken from [16].

## (Un)-certainty in signature gene annotation

Prior to the testing of a gene signature, as a pre-evaluation step, we propose ensuring compatibility between a gene signature and the dataset intended for use. In particular, because of a number of different annotation conventions for genes, compatibility between the genes of a signature - derived from one annotation of the genome, should be able to be mapped to a different annotation of the genome, without significant loss of content or specificity. Because such mappings are generally not bijective, it is critical to ensure that there is reasonable representation of all genes in a signature among the annotation used in a dataset of interest, as this uncertainty can detract from the functional ability of a given gene signature, using publically available tools such as BioMart [5].

## Evaluation of signature gene expression

A critical first step in the evaluation of the validity of a gene signature on a dataset is to ensure that the genes of the signature are expressed at a detectable level across the samples being considered. A gene consistently unexpressed within a gene signature, contributes little to the overall use of the signature as a classifier. Thus *sigQC* evaluates the expression of all genes in the signature, and presents the proportion of samples expressing each gene at supra-threshold level, as well as the proportion of all samples that are not recorded as NA values. The threshold for expression may be user-specified for each dataset, depending also on the platform used. A graphical representation of this in the form of a bar chart and density plot showing the proportion of samples expressing each gene above a particular threshold is returned.

## Evaluation of signature gene variability

In addition to having non-zero expression across a number of samples, signature genes that function well as classifiers should vary across samples. As a result, we propose an evaluation step involving the comparison of the coefficient of variation (a standardised metric of variance) among the genes of the signature, to all genes recorded. This functionality is provided as a visualization of mean versus standard deviation of all genes, and overlaid with the same scatter plot for all signature genes and their associated quantiles for mean and standard deviation.

## Co-correlation of scoring metrics

One need for a gene signature is its ability to be summarised into a ‘scoring’ value to compare across samples. Such a value should encapsulate information from the entire signature, but not be swayed by outliers in the signature genes, which may detract from its performance. To assess this, within *sigQC*, each gene signature score as summarised by the mean, median, and first principal component is taken across the samples, and compared. A high degree of correlation between the metrics gives confidence that the signature score (independent of which metric is used) is providing a reliable value summarising the information of the full signature.

## Effects of data standardisation

A subsequent issue with the application of gene signatures is the effect of data standardisation, as a given signature may be applied on a set of data standardised in a particular way for biomarker discovery purposes, but for application purposes, the data is often re-standardised in a different way. To account for this, we offer that the gene signature score should be compared using non-standardised data and standardised data, to ensure that it ranks samples in a similar way in both cases. In this way, it can be ensured that information carried in the standardisation process will not be lost when using non-standardised data (which we use in clinical application).

## Evaluation of signature compactness

A compact gene signature, often referred to as ‘metagene’, is one that contains genes with high levels of autocorrelation among themselves. To test the level of autocorrelation among signature genes, comparing expression across the dataset, in the *sigQC* package, a heatmap of correlation coefficients is created, which compares the correlation of every gene with every other gene in the signature. Ensuring that all genes act together in a co-ordinated manner ensures that the signature is more likely to have captured a biological response, and that summary scoring metrics will not have significant outlier genes detracting from the other genes of the signature.

## Searching for signature structure

Signature structure can be thought of as an underlying set of components comprising the signature that tend to cluster together in terms of either co-expression, or patient subgroups. Structure can be evaluated using various techniques, here we use hierarchical clustering for its easier visual interpretability with respect to other methods. This initial qualitative assessment is useful to prompt the need for further more advanced analyses of the signature structure. Understanding whether these subcomponents of a signature exist is an important part of evaluating a gene signature, as such clusters may signal either biologically distinct sets of samples within the datasets considered, or may be subgroups of genes that carry redundant information.

## Comparison of multiple signatures and datasets

The *sigQC* package has been designed with an extensible framework, and can be used for the simultaneous evaluation of multiple signatures and datasets at once. A summary plot produced by the package displays a host of metrics summarizing the previous steps on a single radar plot. This methodology facilitates comparison of various metrics of multiple signatures on multiple datasets at once, with a single graphic image. Using this, the quality of various signatures, and the reasons for differences in quality can be rapidly assessed over multiple datasets in a comprehensive manner.

## Evaluation of null distribution of gene signature quality control metrics

Each of the metrics presented on the summary radar plot is computed for a given gene signature on a particular dataset, but to gain a greater understanding of the significance of these values for a given signature, it is critical to consider the underlying null distribution from which each of these statistics arise. Thus, for each dataset and gene signature combination, bootstrap resampling of a random gene signature of the same length as the given signature is computed and presented in boxplot format, for each dataset and for each of the fourteen metrics considered. Immediately, this gives a robust evaluation of the significance of the quality control metrics and how much each differs from its underlying null distribution.

## Comparison with other methods of signature quality control

To our knowledge, no generally adopted methods of gene signature quality control exist in the literature, though some methods have been suggested for specific purposes. For example, validation of significant prognostic ability of a signature is often carried out by resampling random gene lists of the same length as the gene signature, and determining their prognostic ability [17]. This resampling approach is the one adopted by *sigQC* to provide the null distribution for the metrics. Interestingly, such an analysis has also shown that a cutoff of p = 0.05 may be too lenient when aiming to capture specific biology with a gene signature, as many randomly selected gene signatures can also prognosticate with statistical significance [17]. Consensus classification in the face of normalization method uncertainty has also been proposed as an option to robustly evaluate gene signature performance, as done in [6]. These methods solve specific issues related to gene signature validation, but primarily assay the gene signature’s performance as it relates to its final purpose, without consideration for the qualities of the signature genes themselves.

## Limitations

The protocol presented through *sigQC* is limited by the fact that the applicability of a gene signature to a broader setting can never be entirely determined, and so there may be characteristics, intrinsic to a signature or signature types, that enable it to pass all of the proposed quality control measures, without performing well in its intended sense. Such a limitation will almost certainly occur, given the diversity of methodologies of gene signature generation, and to address this, we caution users of this method that it provides a set of conditions which are important to check, but not fully sufficient for the determination of gene signature applicability. Undoubtedly, because of the nature of gene signatures, this limitation will be present regardless of the quality control methodology, as there may always be extreme cases for which such a quality control methodology may not detect a poorly performing signature.

## Materials

### Equipment

#### Hardware

- Personal computer, capable of running R version 3.3.0 or higher

#### Software

- R version ≥ 3.3.0, available to install from https://www.r-project.org/
- *sigQC* package, available to download from https://cran.r-project.org/web/packages/sigQC/index.html

### Equipment setup

#### R software installation

- Download and install the latest version of R from https://www.r-project.org/, or the freely available RStudio from https://www.rstudio.com/.

#### *sigQC* installation

- To install the *sigQC* package, execute the following command in R or RStudio:

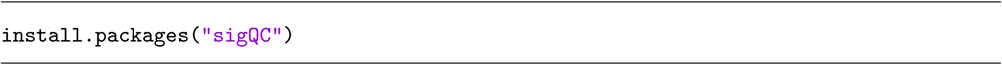

#### Input formats and usage

The primary user-accessible function of *sigQC*, make_all_plots, expects a number of inputs, the format of each of which is defined in Table 1 as well as the package documentation. Further, once installed and with all data loaded into the appropriate variables, use of the package is accomplished with the following commands in R or RStudio:

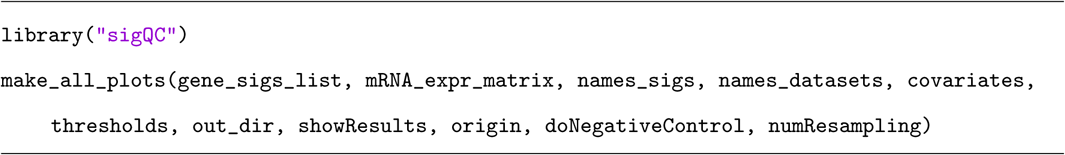

**Table 1:**
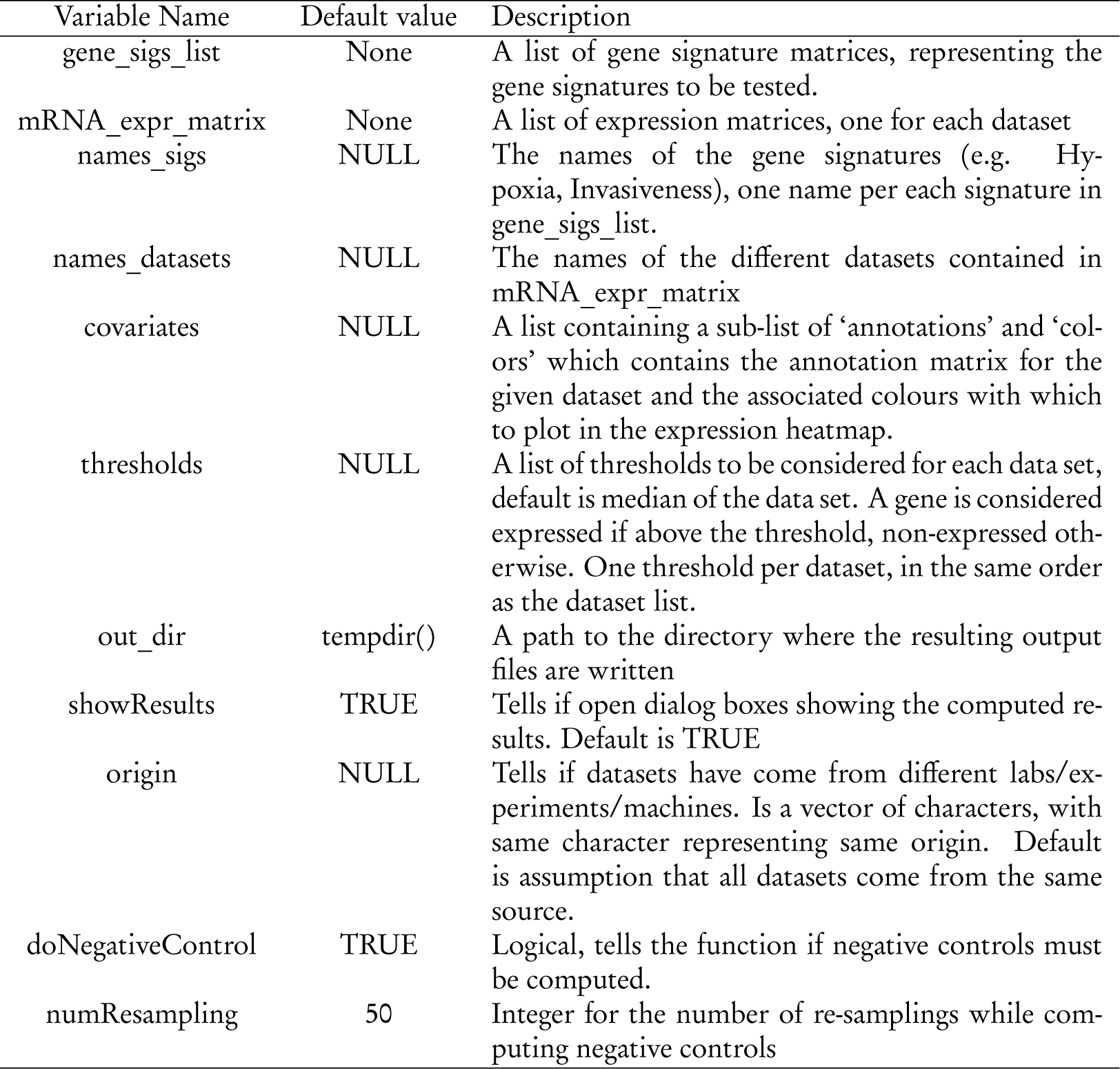
Description of input variables to *sigQC* function make_all_plots().

#### Downloading of sample data and code

Sample randomly generated data and code can be found in the package vignette example that is available for download with the package at https://cran.r-project.org/web/packages/sigQC/index.html

## Procedure

1. Preparation of input data:

- The input data should consist of lists of expression matrices, and should be pre-normalised, and standardised if required. Care should be taken to ensure that genes of interest are present in the dataset and not reported primarily as NA values.
- Additionally, the signatures to be tested must be annotated in a manner consistent with the input data. Furthermore, any specific expression thresholds for expression (other than global median) should be computed, as this is the default the package uses as an expression cutoff.
- Lastly, any additional annotation data to be used alongside the expression heatmaps should be identified, and loaded into the appropriate matrices with color descriptors as specified in the package documentation.
2. Creation of input variables:

- The input data must be loaded into variables in the R environment as specified within the package documentation.
3. Running of *sigQC* package:

- With the input data pre-processed and in the appropriate variables, the principal function of the *sigQC* package can be run, with the following command:

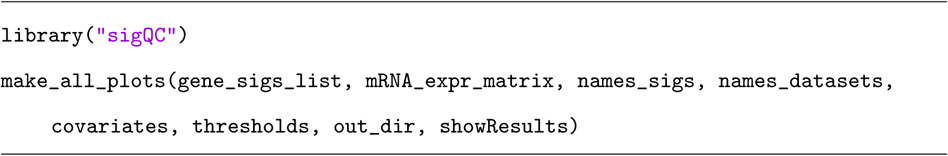
- This produces, in the output directory or graphically displayed directly to the user if desired, a number of plots in PDF files which may be analysed as described in the subsequent steps. The package also creates an output file ‘log.log’ in the output directory, a text file, which summarises the run, and reports any errors that may have occurred if they are not printed to the console. This should be consulted if any issues are encountered in the running of this principal function and for troubleshooting purposes.
4. Analysis of expression:

- From the code and data presented in the Supplementary materials, in steps 4-10 of this procedure, we outline the analysis done for a sample use case of evaluating the suitability of a gene signature on a number of datasets concurrently. We present the use case for the evaluation of a breast cancer metastasis signature and a random set of genes (random gene signature) on the TCGA breast cancer RNA-seq dataset, to show the effects of high and low performing signatures across the quality control metrics. We also present the analysis of output data for the use case of signature transportability across microarray and RNA-seq datasets for the metastasis signature in Box 1.
- First, expression of signature genes should be evaluated across samples in both datasets, and this is done by analysis of the plots sig_expr_*.pdf, as shown in Figure S1A-C. These plots describe the proportion of samples with supra-threshold expression of each signature gene, and the proportion of samples with non-NA values, identifying non-expressed signature components.
5. Analysis of variability:

- The analysis of variability is carried out by loading the file ‘sig_mean_vs_sd.pdf,’ an example of which for sample datasets and gene signatures is shown in Figure 2. These plots describe the mean and standard deviation of expression of all genes reported (in grey) versus all signature genes (in red), with corresponding dashed lines over the plots describing the 10th, 25th, 50th, 75th and 90th percentiles of both mean and standard deviation. This facilitates the easy identification of those signature genes, which are not variable or expressed among the samples, as well as a global evaluation of signature behaviour across samples of a dataset. **Figure 2:**
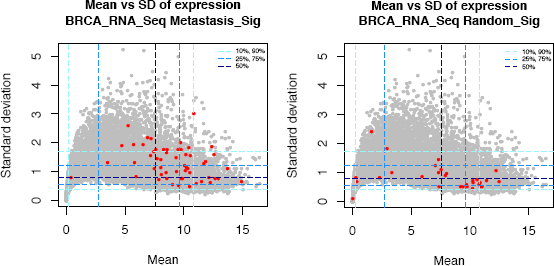
Expression of signature gene expression and variability across datasets for RNA-seq breast cancer for the metastasis signature (left) and a random gene signature (right).
6. Analysis of co-correlation of scoring metrics:

- Loading the files called ‘sig_compare_metrics_*.pdf’ it is possible to analyse the co-correlation of mean, median and first principal component (PCA1) as scoring metrics across the samples for each signature across each dataset, as depicted in Figure 3 Further, also shown in the fourth row of panels of these plots in Figure 3 is a principal components analysis (PCA) scree plot, which describes the proportion of the variance attributable to each principal component, which may reflect whether the first PCA represents a reasonable scoring summary metric for a particular gene signature. **Figure 3:**
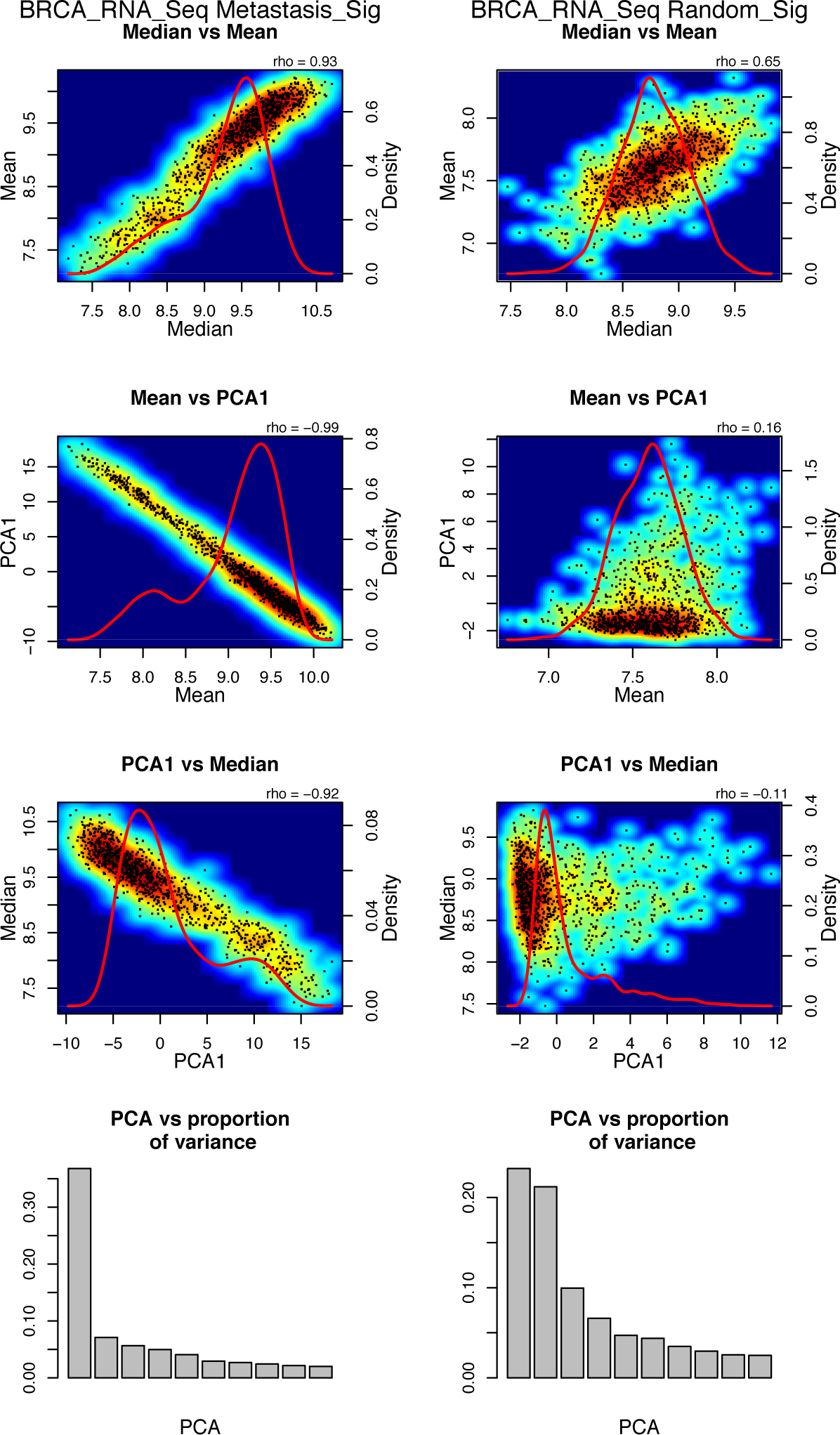
Comparison of scoring metrics for a metastasis gene signature (left) and a set of random genes (right) in the TCGA breast cancer RNA-seq dataset.
7. Analysis of data standardisation effects:

- An analysis of data standardisation effects can be carried out by loading the output file called ‘sig_standardisation_comp.pdf’, an example of which is presented in Figure S2. This plot provides the comparison of median of gene signature expression on the raw data provided versus the median of the gene signature expression on the z-transformed (standardised to zero mean and unit variance) dataset, for each sample in each dataset and each gene signature under consideration.
8. Analysis of signature compactness:

- Loading the files produced in the output directory called ‘sig_autocor_hmaps.pdf’ and ‘sig_autocor_dens.pdf’ provides the plots in heatmap and kernel density estimate plot of the correlation of each signature genes’ expression with the expression of every other signature gene, providing an analysis of signature compactness. We present these plots for the sample data in Figure 4 where it can be seen that as expected, the breast cancer metastasis signature shows a high degree of auto-correlation generally, whereas the random gene signature does not. **Figure 4:**
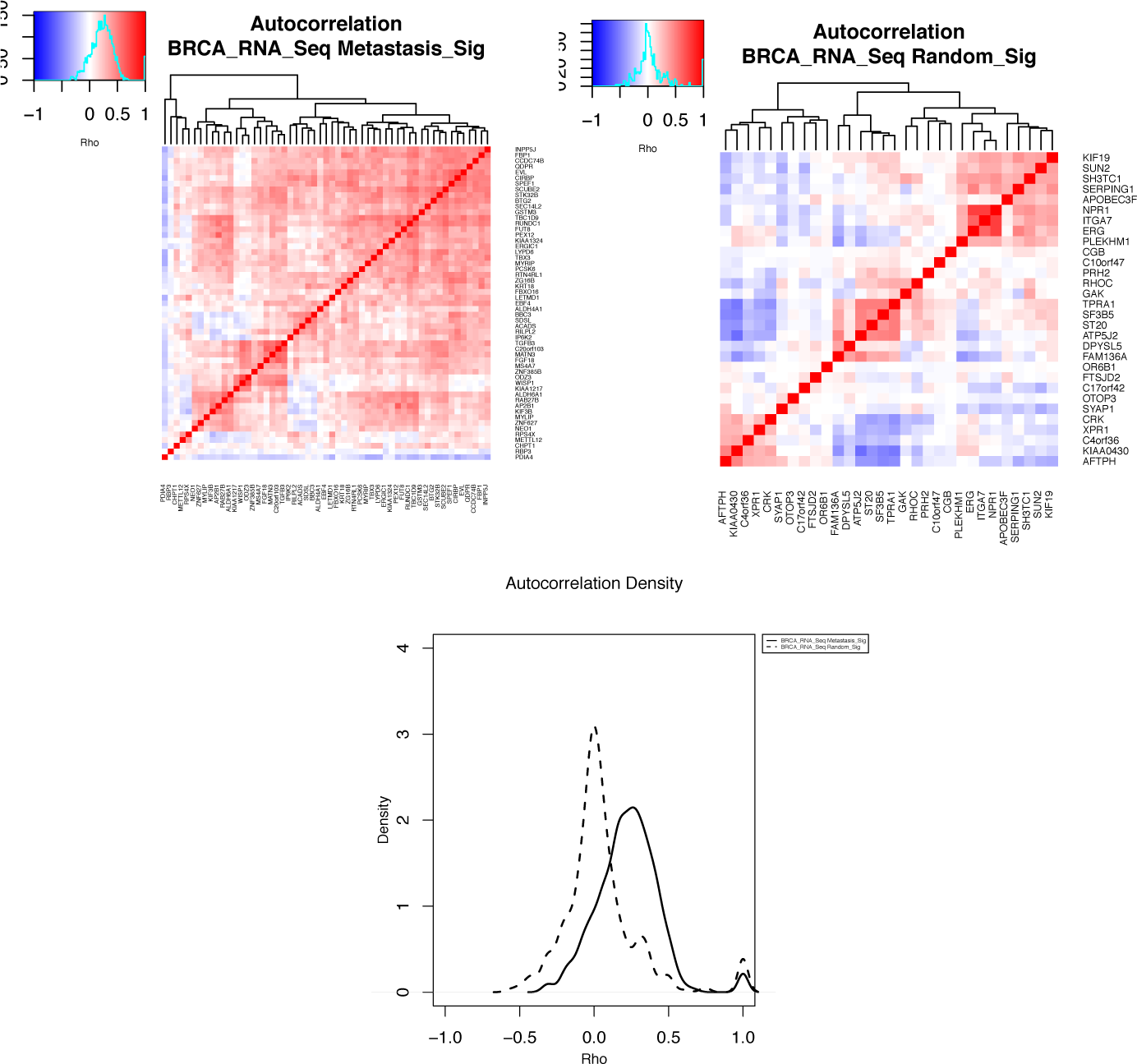
Autocorrelation of signature genes across datasets for a metastasis gene signature (top left) and a random gene signature (top right), with heatmaps represented in density plot form (bottom) for the TCGA breast cancer RNA-seq dataset.
- In addition to the aforementioned plots produced for the analysis of autocorrelation, the files ‘sig_autocor_rankProd_*.pdf’ are produced when there is more than one dataset present for a given gene signature (not shown for this analysis). These plots represent the output of the BioConductor RankProduct package in the evaluation of signature genes whose median autocorrelation with all other genes consistently ranks low with the other signature genes. This facilitates the process for finding genes with consistent poor autocorrelation with the other genes of the gene signature, particularly when refining a given signature for optimal performance across a number of different datasets (e.g. multiple clinical cohorts, or clinical data and cell line data).
9. Analysis of signature structure:

- Signature structure is evaluated by the consideration of a number of plots created by the *sigQC* package. Firstly, signature structure is sought to be evaluated by hierarchical clustering on the provided expression values of the signature elements over all samples, in conjunction with annotations for the samples, if they are provided. These plots are present in the output directory and are named ‘sig_eval_struct_clustering_*.pdf’, which are clustered based on each dataset in turn, and run over each signature and each dataset present. An example of such a plot is shown in Figure 5, where the different expression profiles of the random gene signature and the metastasis gene signature across patients can be seen. **Figure 5:**
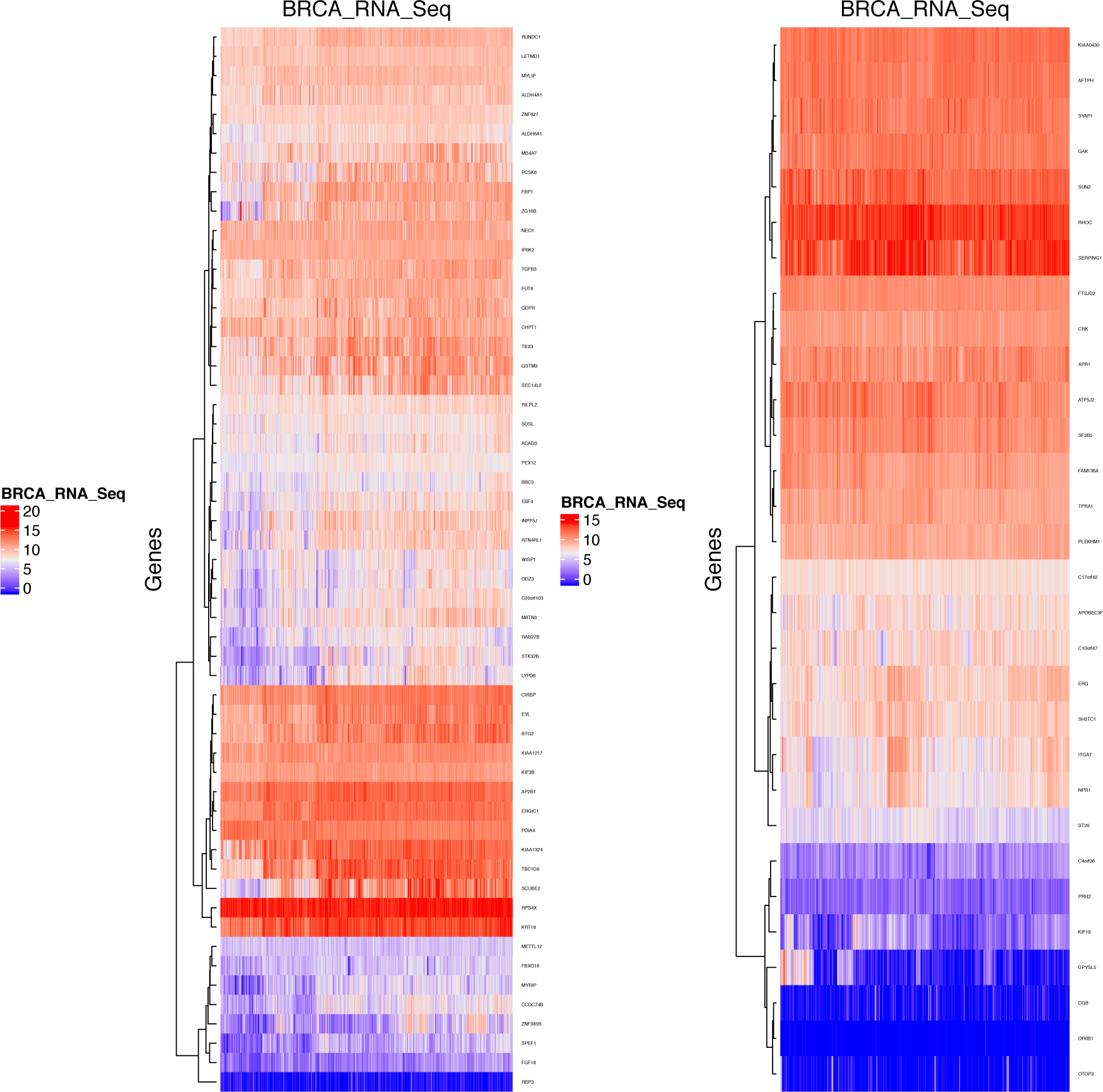
Hierarchical clustering of signature gene expression for the metastasis signature (left) and the random signature (right) over the TCGA breast cancer dataset.
- In addition to hierarchical clustering on patient samples and signature elements, in the output directory of *sigQC*, biclustering results can be found, which describe the output if biclusters of sample groups and signature elements are found (not shown for this analysis). The particular details of the biclustering outputs and algorithm used may be found in the R Package documentation. If no biclusters are found, the files termed ‘sig_eval_bivariate_clustering.pdf’ show blank plots only, as had occurred with the sample datasets and signatures presented.
10. Optional: Comparison of multiple signatures:

- The file produced, entitled ‘sig_radarplot.pdf’ describes each signature applied to each dataset in a holistic, radar chart format. This plot evaluates the gene signature across a number of metrics, many of which are summary metrics for those in steps 4-9 of this procedure, and these are described in detail in the Supplementary Table S1. A sample of this plot, for the metastasis gene signature and the random gene signature on the TCGA breast cancer data set is shown Figure 6. **Figure 6:**
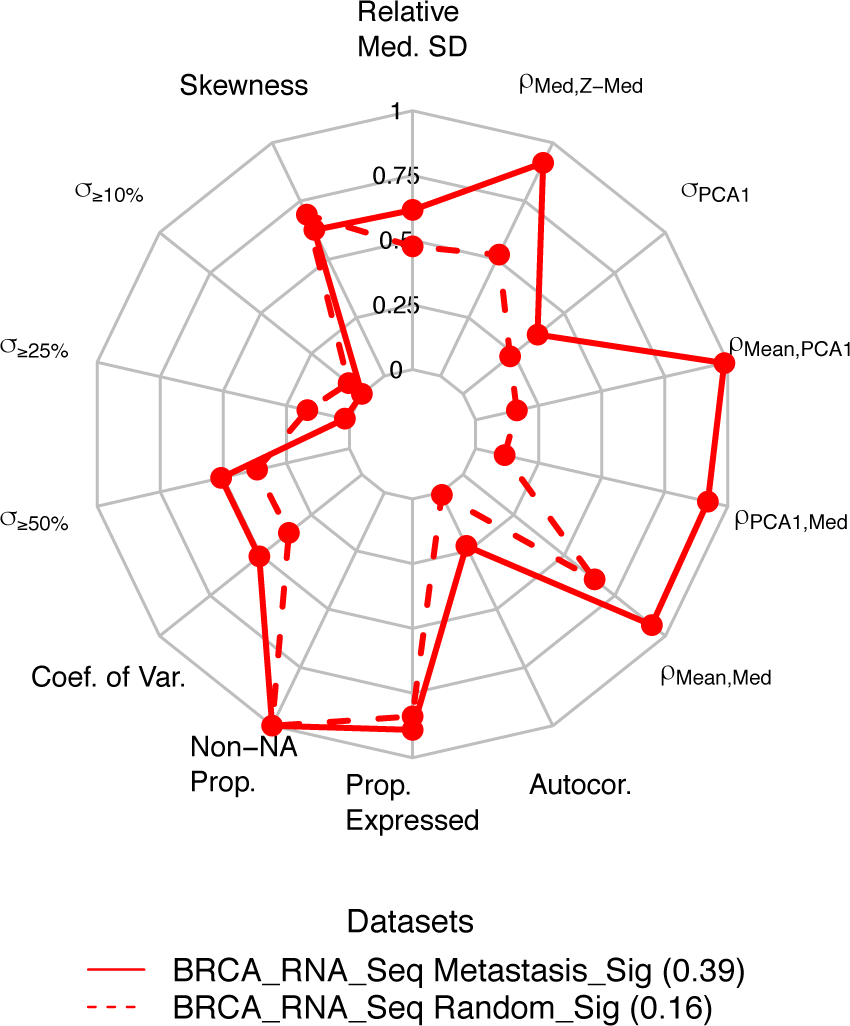
Radar plot showing summary of gene signature quality control metrics for a metastasis signature (solid line) and a random set of genes (dashed line), as calculated for the TCGA breast cancer RNA-seq dataset.
11. Optional: Analysis of null distributions of QC metrics (Timing: several minutes-hours):

- The file produced, entitled boxplot_metrics.pdf, in the negative_control subfolder of the results output shows the distributions of each of the fourteen summary metrics as reported in the radar plot for each signature and dataset combination. These distributions are generated for the number of repeats as specified by the input parameter numResampling, with default set to 50. The values for the gene signature and dataset combination in question are shown in red overlaid with the other points in grey, giving a sense of significance of each of the metrics, as shown in Figure 7. From this figure, it can be seen that for the breast cancer metastasis signature on this dataset, the metrics evaluated show high significance for the signature genes, as compared to a random set of genes of the same length, whereas the same significance is not seen for the randomly chosen gene signature. **Figure 7:**
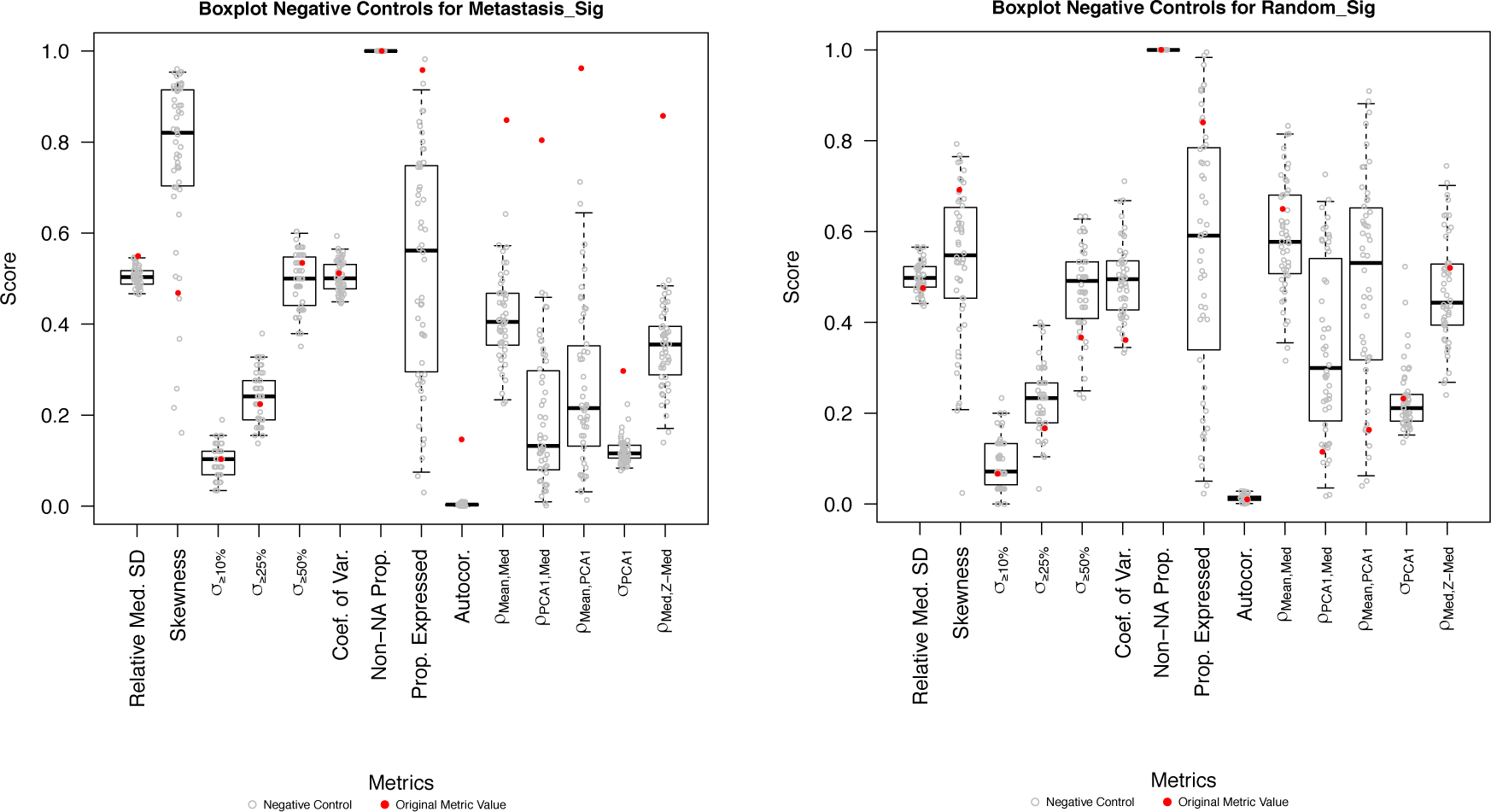
Box and scatter plots depicting the null distributions of each of the metrics measured on the radar plot for the metastasis signature (left) and a random signature (right), for *N* = 50 resampling runs using random gene signatures of the same length.
12. Optional: Analysis of raw data:

- The package produces tables of raw data, in tab-delimited text format, output into subfolders in the output directory for ease of re-analysis by the user. In particular, the *sigQC* package produces subfolders for the table of mean expression and standard deviation of expression for all signature elements, tables of the mean, median, and first PCA of each sample, tables of the median and z-transformed median for each sample, autocorrelation matrices for all signature elements, tables of proportion of expression above threshold and proportion of NA expression for all signature elements, as well as the table of values plotted for each signature and dataset in the summary radar plot. We provide these raw data tables so that they may then be reloaded into the user’s analysis pipeline of choice for re-plotting or re-analysis.

## Timing

The timing of *sigQC* functions varies, depending on the number of datasets and signatures analysed, from few minutes (for the examples shown here) to hours (for concomitant analysis of several datasets and signatures, and high number of replicate resampling).

## Troubleshooting

### Installation

Issues may be experienced if the ‘ImageMagick’ dependency is not installed on the user’s system (particularly for Windows systems). To install this dependency, please follow directions at: http://imagemagick.org/script/download.php.

### Step 3

Issues may be experienced with input data not conforming to the format required by *sigQC*. If this occurs, the package will alert the user with an error message describing the nature of the discrepancy. For example, common errors may include the following:

- Gene signatures must be formatted as a list of matrices, of dimension *k* rows by 1 column, for a signature of length *k* genes. Inputting a single list as a vector will cause an error to the program.
- Datasets must also be formatted as lists of matrices, such that genes are the rownames of the dataset, and samples are organised by columns of the dataset.
- Gene signatures and datasets must be annotated in the same way, as if the names of the genes of a signature are not found in a dataset, the computation will not continue.
- Care must be taken to ensure that NA valued genes are removed as optimally as possible, as if there are too many values in the expression matrix for the gene signature are NA, calculations dependent upon singular value decomposition (e.g. principal components analysis) cannot be carried out.

## Anticipated results

Here, we provide an explanation of the results generated in the example described above. In particular, we describe the figures produced in steps 4-11 of the above procedure.

### Analysis of expression and variability (Steps 4-5)

In the case of a signature performing well on a given dataset, the genes of the signature will be highly expressed and highly variable, as evidenced by the plots of expression and variability in Figures 2 and S1. As shown in Figure 2, the red dots, corresponding to the genes of the signatures are enriched higher-expression and higher-variability regions of the plot for the metastasis signature, as compared to the random gene signature.

### Analysis of scoring metrics and standardisation (Steps 6-7)

The next steps in the protocol are the evaluation of the correlation between different scoring metrics, and whether standardisation preserves scoring metrics’ rank within the dataset. In the case of a well-performing signature on a dataset, as evidenced by the case of the metastasis signature, as seen in Figures 3 and S2, each of the scoring metrics is correlated with the other, as well as the median metric between standardised and un-standardised data. This is not seen as such for the random gene signature, as might be expected.

### Analysis of autocorrelation (Step 8)

Within a well-performing gene signature, each of the genes of the signature should be acting within the same way within the signature; that is, each gene should be increasing or decreasing in expression concordant with the others of the signature. This is quantified by the autocorrelation of the gene signature, as shown in Figure 4, and it can be seen that for a well-performing signature, the metastasis signature, there is a significant autocorrelation between each of the genes, which is not seen for the random gene signature.

### Analysis of signature structure (Step 9)

For a signature expected to perform well on a full dataset, each of the genes should have an expression profile similar to itself across all patients; that is, there should not be necessarily subgroups of patients with markedly different expressions of the genes of the signature discordant with other genes of the signature (i.e. all genes should act in a similar manner to capture a biological phenomenon). This is evidenced by the lack of biclusters and obvious visual subclusters of patients for both signatures considered in this example in Figure 5.

### Global analysis of metrics and their significance (Steps 10-11)

In order to effectively compare signatures across a range of metrics, we designed the radar plot depicted in Figure 6, to quickly show, on a scaled plot, the various means of comparing statistical properties of gene signatures across datasets. As can be seen, over nearly all metrics, the metastasis gene signature outperforms the random gene signature, as might be expected. However, to fully appreciate the magnitude of these differences, an understanding of the null distribution of each of these metrics is required, which is shown in Figure 7, from Step 11. This shows that over nearly every metric, the well-performing gene signature for breast cancer metastasis is highly significant, whereas the random gene signature does not show significance across many of the metrics considered, thereby facilitating the rapid identification of a well-performing versus a poor-performing gene signature.

#### Box 1

##### Example evaluation of signature translatability cross-platform

As an example to highlight the utility of *sigQC* in determining the translatability of a signature across different sequencing platforms, we consider here the outputs of the package in comparing cross-platform performance. In particular, we consider the same breast cancer metastasis signature [16] taken from MSigDb, on each of an RNA-seq dataset (TCGA), and a microarray generated dataset (GEO Series GSE3494). An initial step is to generate a signature annotated for each of the platforms, which is done through the use of BioMart, enabling the conversion of gene symbols into Affymetrix U133A probe IDs for use with the microarray dataset. Subsequently, running *sigQC* on each of these datasets and converted signatures individually gives the underlying data needed to generate the following radar plots, which can be used, in conjunction with the plots of negative control, to determine both the differences and the significance of these differences of the metrics reported by the radar plot. In this example, we observe that there is a high concordance between the outputs of the radar plots in both case, as well as high significance of many of the metrics in both cases, suggesting that this signature is highly applicable cross-platforms.

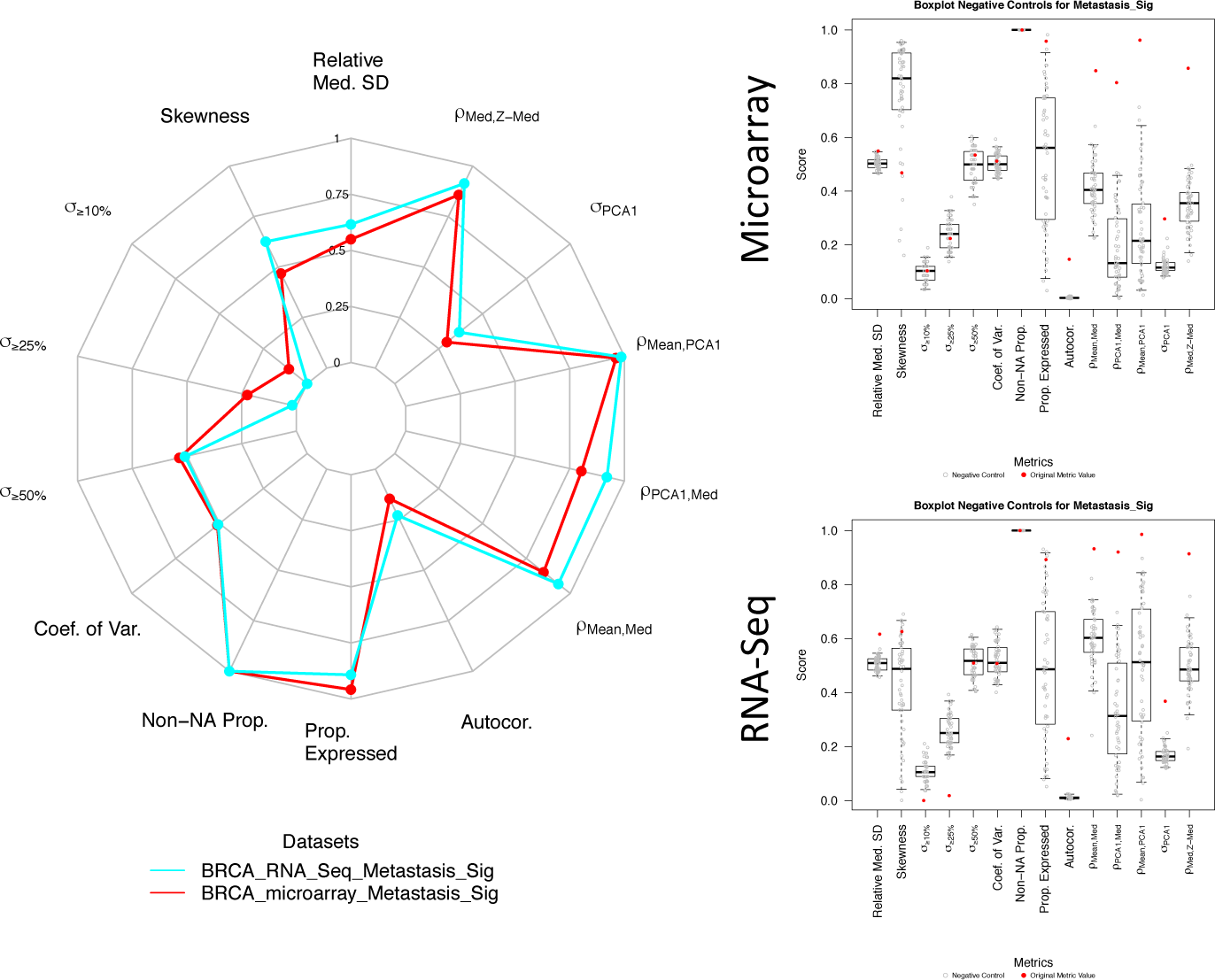

## Acknowledgments

This work was funded by Cancer Research UK (F.M.B., A.L.H., A.B., W-C.C., A.D.) and the Medical Research Council (T.M., E.D.). The authors are also grateful for the support of the Clarendon Fund to A.D.

## Author contributions

F.M.B. conceived the idea and designed the study. A.D., A.B., W-C.C., J.S. and F.M.B. contributed statistics and data visualization. A.D. performed analyses. A.D., A.B. and W-C.C. wrote and debugged code. A.B. and F.M.B. supervised the implementation. A.L.H., T.M., E.D. and C.W. contributed application cases and interpretation of data. A.D., A.B., J.S. and F.M.B. wrote the manuscript with contribution from all other authors.

## Supplementary information

### S1: *sigQC* availability

The *sigQC* package has been made available for download from CRAN at https://cran.r-project.org/package=sigQC, and can be cited through this publication.

### S2: Radar plot metrics

In Table S1 we provide a description of the metrics plotted on the arms of the radar plot.

**Table S1:**
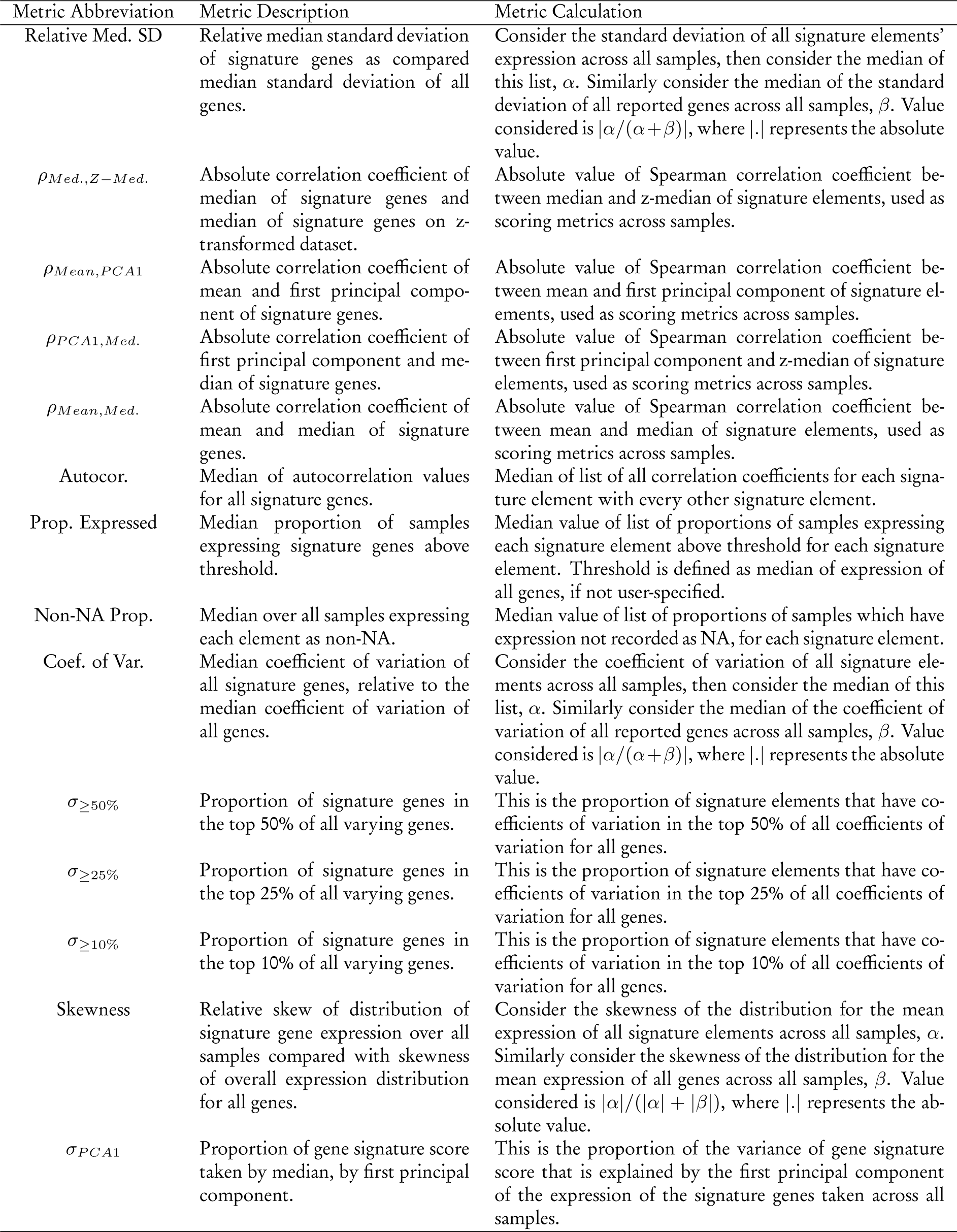
**Description of metrics defining components of summary radar plot.**

### S3: Pseudocode for radar plot metrics

We define an m-dimensional array, *e* = [*e*_1_, …, *e_m_*] as the gene expression data relative to a single sample, such that *e_k_* is the expression value of gene k in the given sample. In this way, we may define the full dataset as the bi-dimensional matrix*E* = [*e*_1_, …, *e_n_*], where *n* is the number of samples and *e_ij_* is the expression value of gene *i* in the *j*-th sample. Similarly, we denote by *E* = [*e*_1_, …, *e_m_*]^*t*^ the same matrix, where *e_k_* is an n-dimensional array containing the expression data of a single gene across all *n* samples and (*·*)^*t*^ indicates the transpose of a matrix. Finally, we denote by *R* = [*r*_1_, …, *r_n_*] the reduced gene expression matrix containing only the expression of the genes included in the assessed signature so that *r_k_* = [*r*_1_, …, *r_l_*], where *l* ≤ *m*.

#### Ratio of Med. SD

1. Compute the standard deviation (*σ*_1_) of each signature gene across all samples
2. Denote by α the median of the standard deviations
3. For every gene, compute the standard deviation (*σ*_2_) across all samples
4. Denote by β the median of the standard deviations
5. Return the absolute value of *α*/(*α*+*β*)

##### Pseudocode

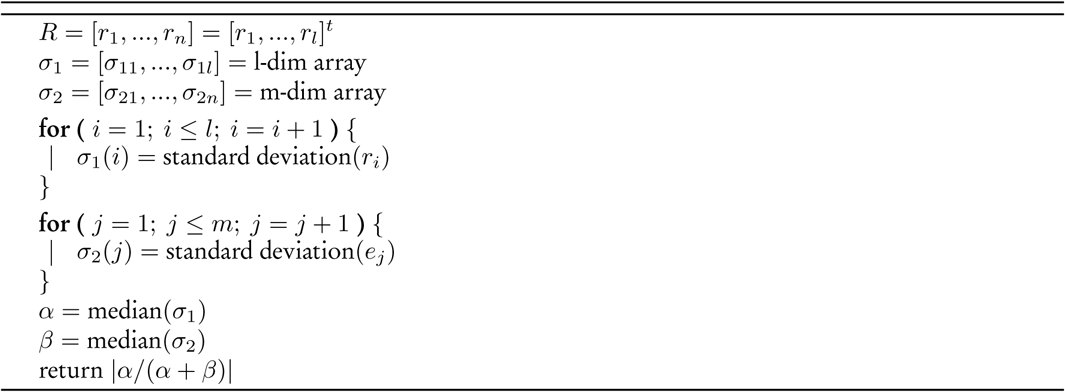

#### Med., Z-Med. Score Cor

1. Compute the median of each signature gene across all samples
2. Normalise the input matrix using the z score
3. Compute the median of each signature gene in the normalised matrix across all samples
4. Compute the Spearman correlation between the 2 median arrays
5. Return the absolute value of the Spearman correlation coefficient

##### Pseudocode

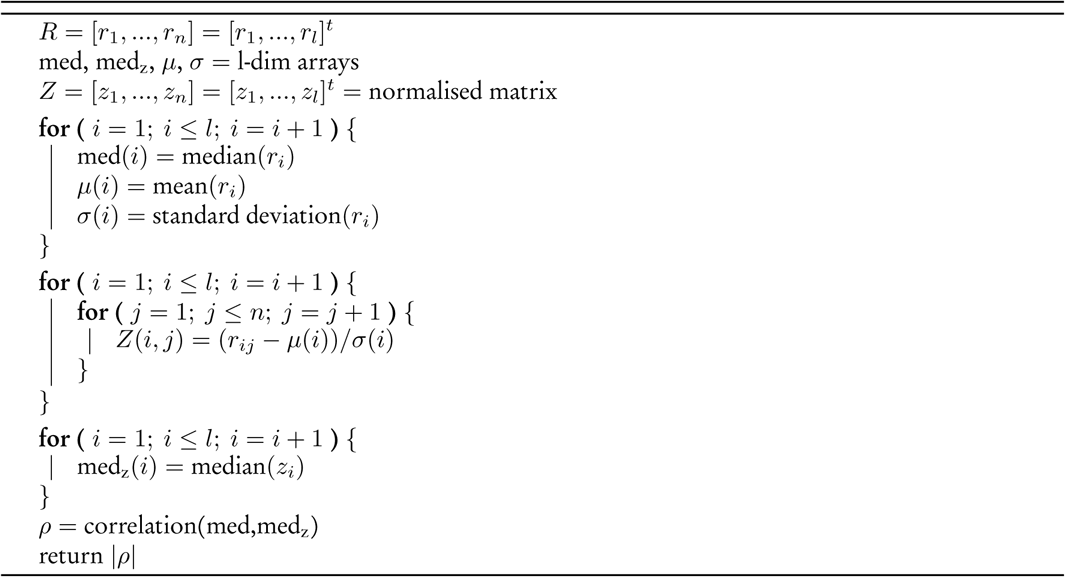

#### Mean, PCA1 Score Cor

1. Compute the mean of each signature gene across all samples
2. Compute the first principal component (PCA1) of each signature gene across all samples
3. Compute the Spearman correlation between the mean and PCA1 arrays
4. Return the absolute value of the Spearman correlation coefficient

##### Pseudocode

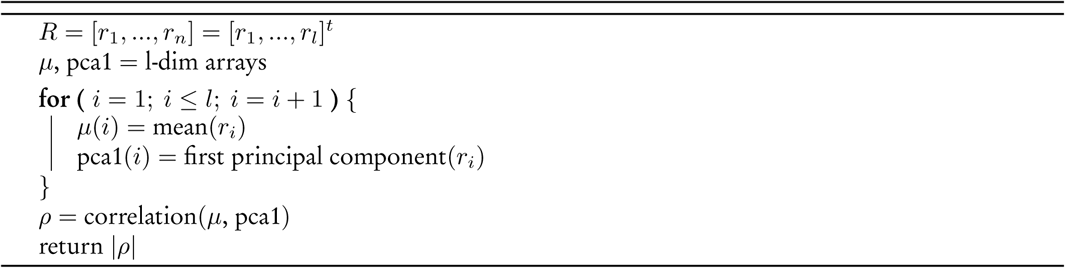

#### PCA1, Z-Med. Score Cor

1. Compute the first principal component (PCA1) of each gene across all samples
2. Normalise the input matrix using the z score
3. Compute the median for each signature gene across all samples in the normalised matrix
4. Compute the Spearman correlation between the PCA1 and median arrays
5. Return the absolute value of the Spearman correlation coefficient

##### Pseudocode

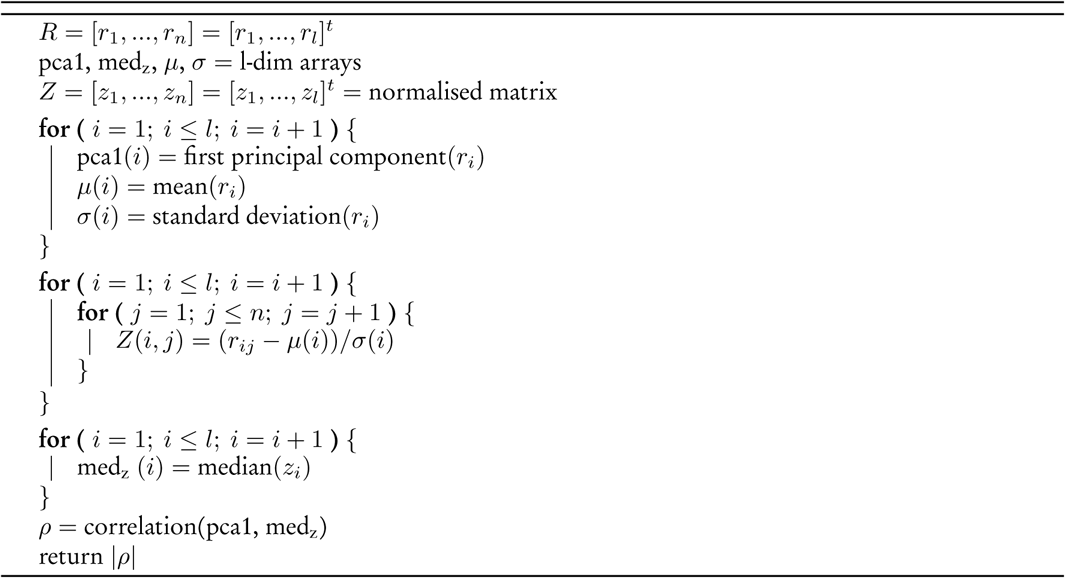

#### Mean, Med. Score Cor

1. Compute the mean of the signature genes for each sample
2. Compute the median of the signature genes for each sample
3. Compute the Spearman correlation of the mean and median arrays
4. kReturn the absolute value of the Spearman correlation coefficient

##### Pseudocode

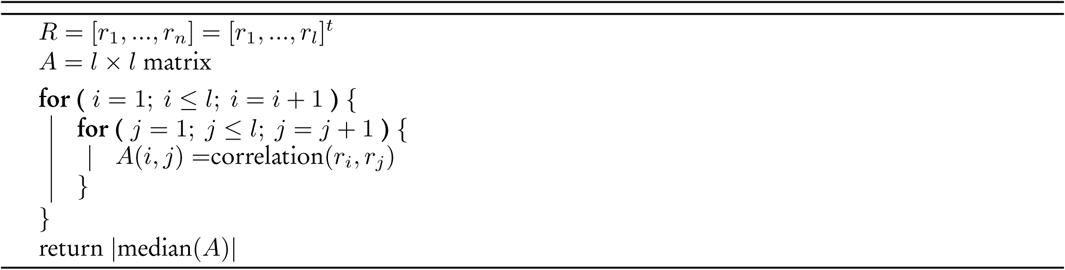

#### Med. Autocor

1. Compute the autocorrelation of the reduced gene expression matrix
2. Return the absolute value of median of all correlations coefficients

##### Pseudocode

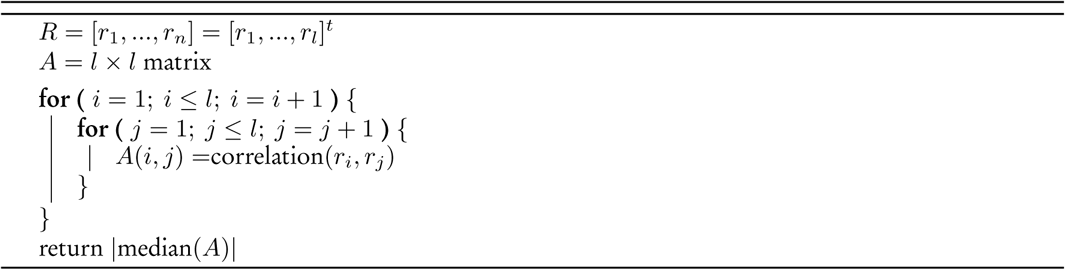

#### Med. Prop. Expressed

1. Compute the median of the dataset
2. For each gene, check if expression is greater than median
3. For each gene, count the proportion over all samples
4. Return the median over the array of proportions

##### Pseudocode

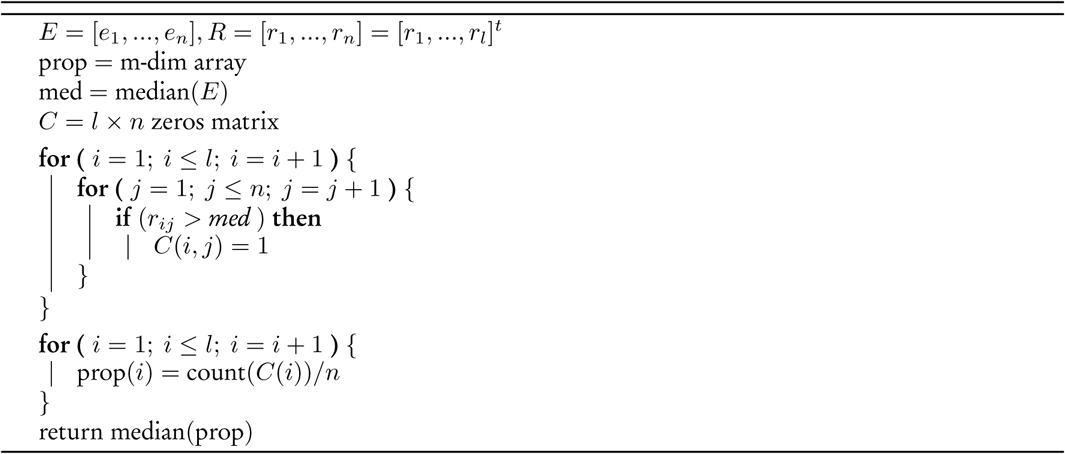

#### Med. non-NA Prop

1. Count the number of times each gene in the signature is expressed over all samples
2. For each gene, compute the expression proportion over all samples
3. Return the median over the array of proportions

##### Pseudocode

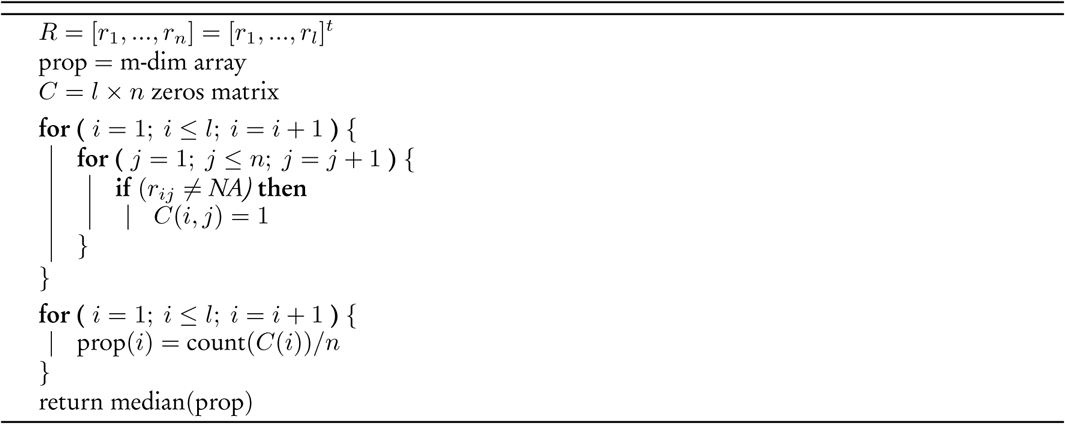

#### Coef. of Var. Ratio

1. Compute the standard deviation (*σ*) for each signature gene across all samples
2. Compute the mean (*μ*) for each gene across all samples
3. Compute the coefficient of variation (*c*_*v*1_ = *σ/μ*) for each signature gene across all samples
4. Denote by α the median of the coefficients of variation
5. For each gene, compute the coefficient of variation (*c*_*v*2_) across all signature genes
6. Denote by β the median of all *c*_*v*2_
7. Return the absolute value of *α*/(*α*+*β*

##### Pseudocode

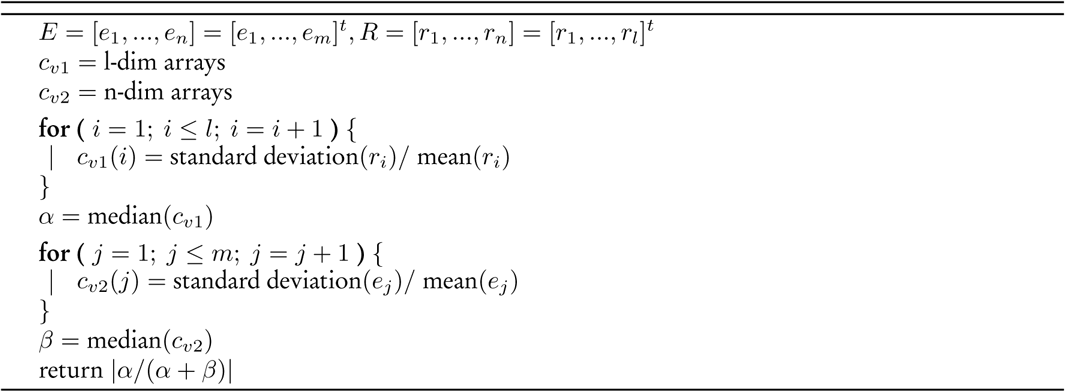

#### Prop in top 50% var

1. Compute the standard deviation (*σ*) for each gene across all samples
2. Compute the mean (*μ*) for each gene across all samples
3. Compute the coefficient of variation (*c*_*v*_ = *σ*/*μ*) for each gene across all samples
4. Rank the *c_v_*
5. Return the proportion of signature genes with *c_v_* in the top 50% of the rank

##### Pseudocode

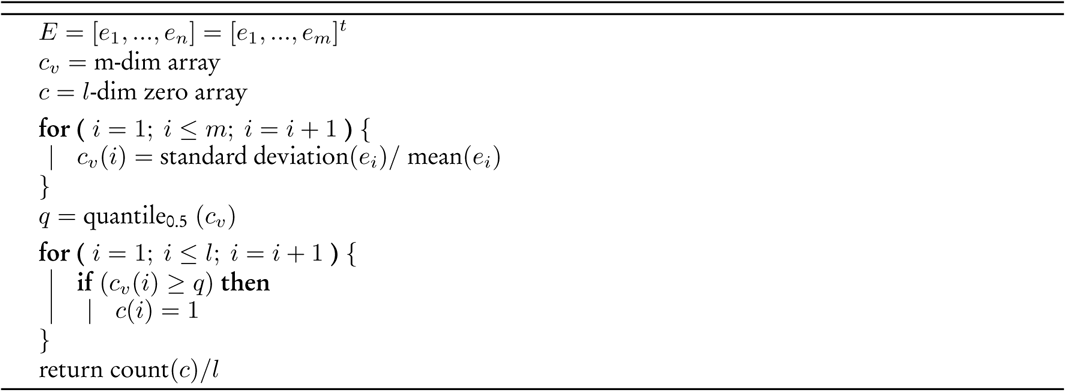

#### Prop in top 25% var

1. Compute the standard deviation (*σ*) for each gene across all samples
2. Compute the mean (*μ*) for each gene across all samples
3. Compute the coefficient of variation (*c*_*v*_ = *σ*/*μ*) for each gene across all samples
4. Rank the *c_v_*
5. Return the proportion of signature genes with *c_v_* in the top 25% of the rank

##### Pseudocode

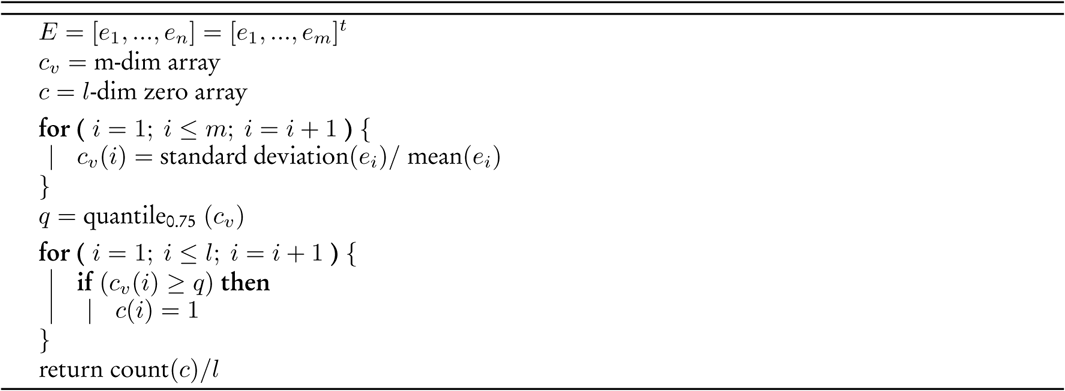

#### Prop in top 10% var

1. Compute the standard deviation (*σ*) for each gene across all samples
2. Compute the mean (*μ*) for each gene across all samples
3. Compute the coefficient of variation (*c_v_* = *σ*/*μ*) for each gene across all samples
4. Rank the *c_v_*
5. Return the proportion of signature genes with *c_v_* in the top 10% of the rank

##### Pseudocode

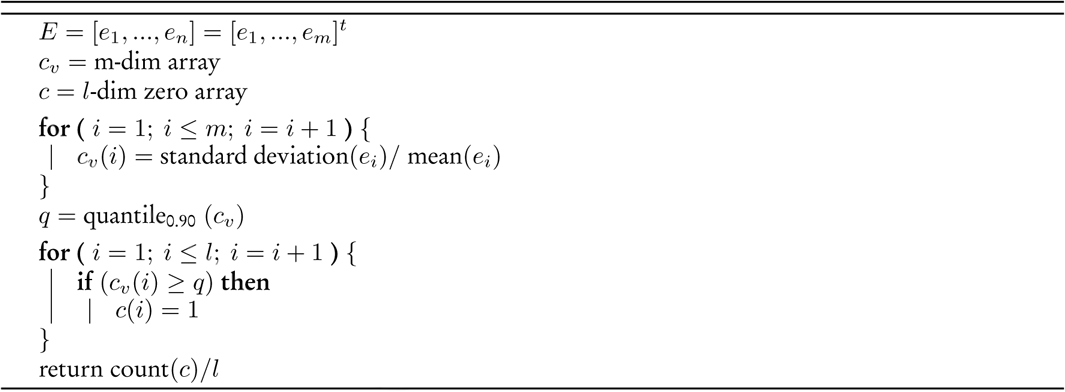

#### Skew Ratio

1. Compute the skewness (*α*) of mean of each signature gene, across all samples
2. Compute the skewness (*β*) of mean of each gene, across all samples
3. Return |*α*|/(|*α* + *β*|)

##### Pseudocode

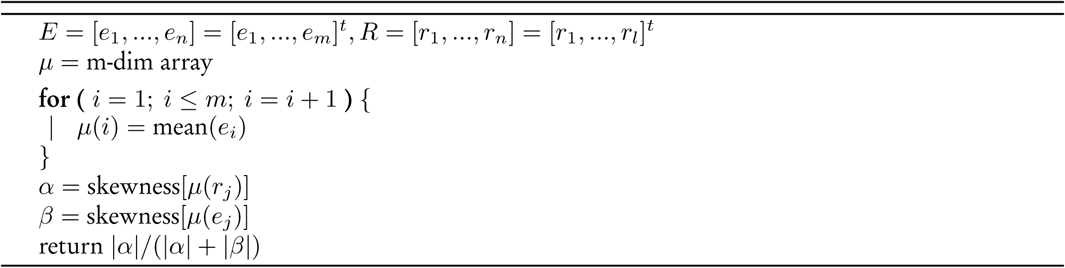

#### Prop Var by PCA1

1. Compute the principal component of every signature gene across all samples
2. Return proportion of variance explained by first principal component

##### Pseudocode

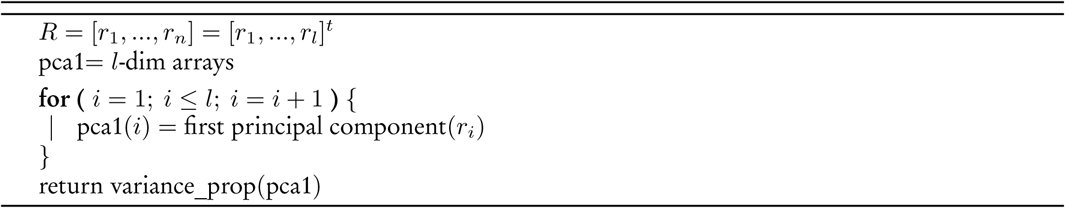

## S4: Supplementary figures

**Figure S1:**
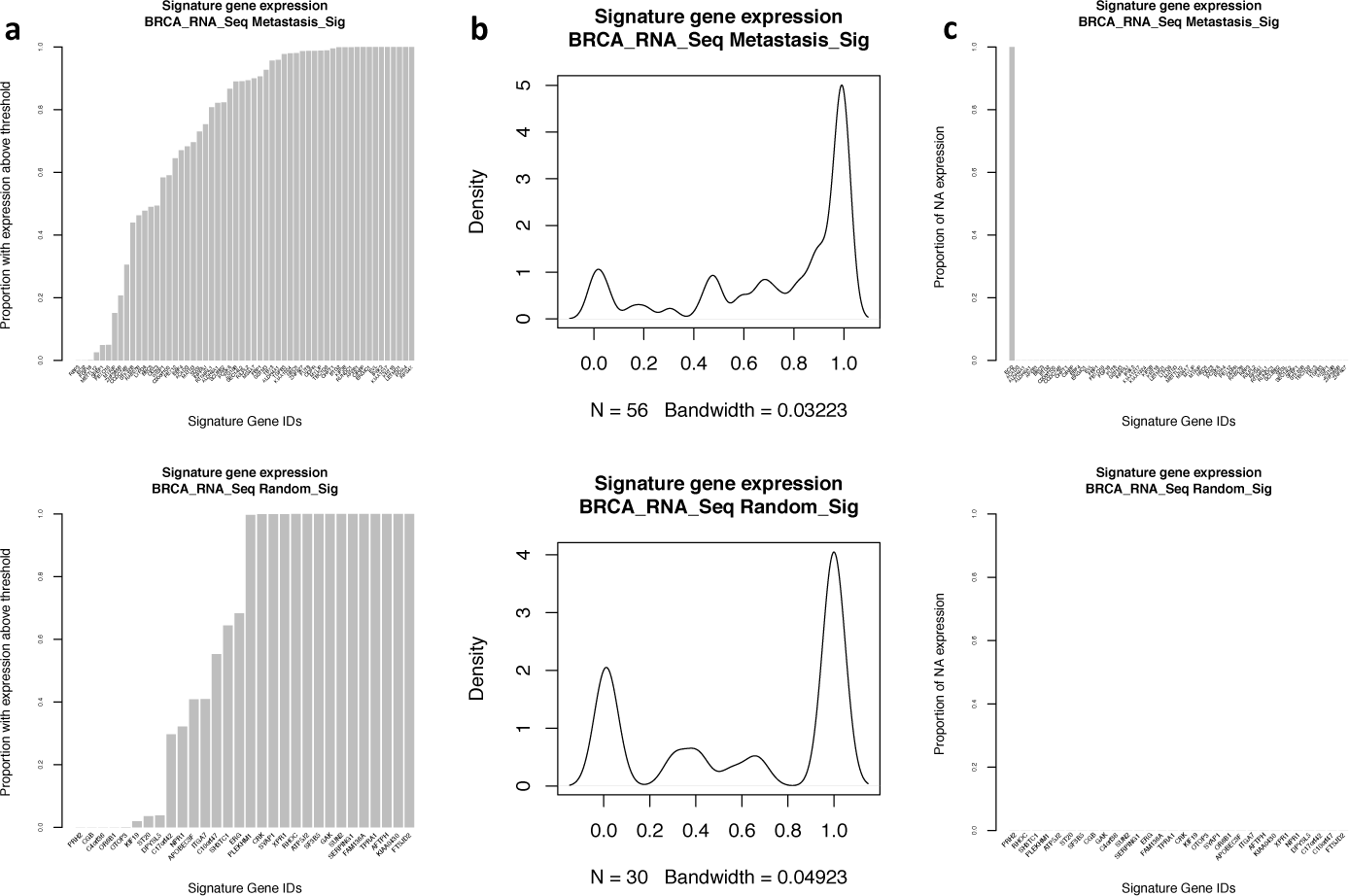
Expression of signature genes across the TCGA breast cancer RNA-seq dataset for the metastasis gene signature (top) and a random set of genes (bottom), shown as (a) a barplot for the proportion of samples expressing a gene above the median, (b) a density plot showing the same information as the barplots in (a), and (c) a plot of the proportion of samples showing NA expression for each of the genes of the signature.

**Figure S2:**
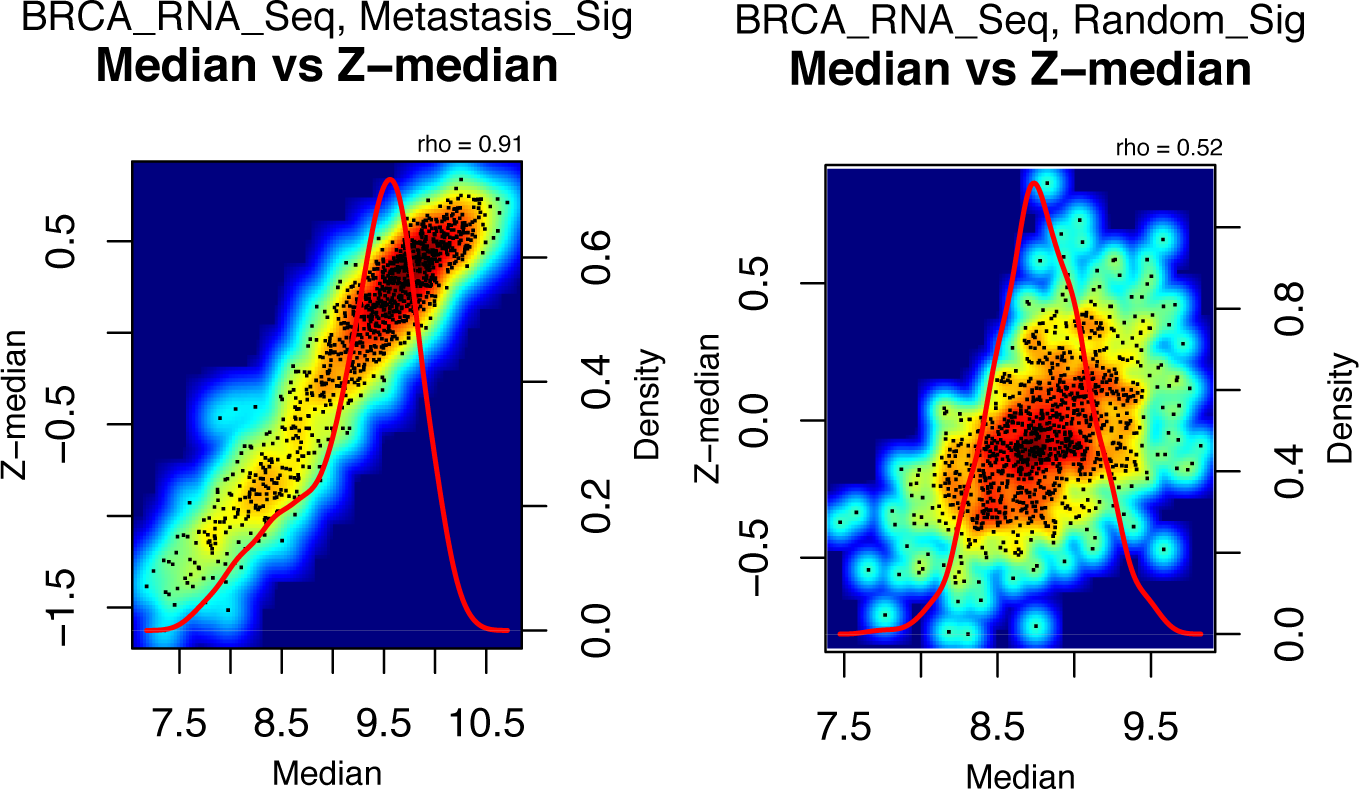
Comparison of median and z-transformed median of signature gene expression across the RNA-seq breast cancer dataset for the metastasis gene signature (left) and the random set of genes (right).

